# The DNA event horizon in the Guaymas Basin subsurface biosphere: technical advances and re-defined limits in bulk extractions of nucleic acids from deep marine sediments

**DOI:** 10.1101/2024.04.15.589529

**Authors:** Gustavo A. Ramírez, Paraskevi Mara, David Beaudoin, Diana Bojanova, John E. Hinkle, Brewster Kingham, Virginia P. Edgcomb, Yuki Morono, Andreas Teske

**Affiliations:** Dept. of Biological Sciences, California State University Los Angeles, Los Angeles, CA, USA; Dept. of Geology & Geophysics, Woods Hole Oceanographic Institution, Woods Hole, MA, USA; Dept. of Earth Sciences, University of Southern California, Los Angeles, CA, USA; Dept. of Earth, Marine and Environmental Sciences, University of North Carolina at Chapel Hill, Chapel Hill, NC, USA; DNA sequencing & Genotypic Center, University of Delaware, Newark, DE, USA; Kochi Institute for Core Sample Research, Institute for Extra-cutting-edge Science and Technology Avantgarde Research (X-STAR), Japan Agency for Marine-Earth Science and Technology (JAMSTEC), Monobe, Nankoku, Kochi, Japan

## Abstract

We compiled DNA and RNA isolation protocols for sediment bulk extraction and their yields from Guaymas Basin subsurface sediments, and evaluated their sensitivity for metagenomic and amplicon analyses of subsurface microbial communities. Guaymas Basin sediments present a challenge for DNA and RNA recovery due to high concentrations of hydrocarbons, steep thermal gradients and rapidly declining cell numbers downcore. Metagenomic library construction and sequencing was possible from as little as 0.2 to 0.5 ng DNA/cm^3^ sediment; PCR amplification of 16S rRNA genes required in most cases approx. 1-2 ng DNA/cm^3^ sediment. At in-situ temperatures of 50 to 60°C, decreasing DNA recovery leads to increasingly uncertain “hit or miss” outcomes and to failures for metagenomic and amplicon analyses. DNA concentration profiles show that, even before these hot temperatures are reached, relatively moderate temperatures have a major effect on microbial abundance and DNA yield. Comparison with cell count profiles shows that hydrothermal influence is reducing downcore cell densities by multiple orders of magnitude faster compared to non-hydrothermal sediments; this effect is also visible at relatively moderate temperatures. To an even greater degree than DNA, RNA recovery is highly sensitive to downcore increasing temperatures and decreasing cell numbers, and worked best for microbial communities in cool, relatively shallow subsurface sediments.

## Introduction

Cultivation-independent sequencing studies of microbial communities in the deep subsurface require extraction of nucleic acids in sufficient quantity and quality for subsequent high-throughput sequencing. Here we examine the outcome of DNA and RNA extractions for deep subsurface sediments from Guaymas Basin, a sedimented, hydrothermally active spreading center with steep thermal gradients and high heat flow in the axial troughs and flanking regions (Neumann et al., 2023). We focus on Guaymas Basin sediments because among deep-sea sediments they are some of the most challenging sediments to extract nucleic acids from due to their high content of organic material, including hydrocarbons that can interfere with the activities of enzymes and other reagents in extraction protocols. The observations we make about downcore DNA and RNA recovery will not only be useful for the community of scientists interested in the Guaymas Basin, but also for the broader community of scientists who study the sedimented subsurface biosphere.

Eight different sites with contrasting thermal and geochemical regimes were drilled during IODP expedition 385 (Teske et al., 2021a). The positions of these drilling sites generally followed a northwest-to-southeast transect across the northern Guaymas Basin flanking regions and the axial trough. Two neighboring sites (U1545 and U1546) on the northwestern end of Guaymas Basin (Teske et al., 2021b; Teske et al., 2021c) essentially differ by the presence of a massive, thermally equilibrated sill between 350 to 430 mbsf at Site U1546 (Lizarralde et al., 2023). Two drilling sites (U1547, U1548) target the hydrothermally active Ringvent area, approximately 28 km northwest of the spreading center (Teske et al., 2019), where a shallow, recently emplaced hot sill creates steep thermal gradients and drives hydrothermal circulation (Teske et al., 2021d). Drilling Site U1549 (Teske et al., 2021e) explores the periphery of an off-axis methane cold seep, Octopus Mound, located ∼9.5 km northwest of the northern axial graben (Teske et al., 2021f). Of all these sites, the two Ringvent sites have the steepest thermal gradients (between 506 and 958°C per km) and the highest heatflow values (between 516 and 929 mW per m^2^) (Neumann et al., 2023). This distinct thermal regime is caused by a recently emplaced, shallow volcanic sill that is driving local hydrothermal circulation (Teske et al., 2019). Initial shipboard cell counts performed on the RV JOIDES *Resolution* indicate rapidly decreasing cell numbers with depth in all sites, starting above 10^9^ cells per cm^3^ in surficial sediments, but decreasing towards 10^6^ cells per cm^3^, and lower (Teske et al., 2021a). The downcore decrease in cell numbers is steeper at Ringvent compared to the other sites; values near 10^6^ cells per cm^3^ are reached around 60 meters below sea floor (mbsf) at Ringvent, whereas comparable cell densities at the northwestern sites U1545 and U1546 are reached only around 150 mbsf (Teske et al., 2021a). In first approximation, this steep downcore decline in cell densities appears to be linked to the strong thermal gradients in the Guaymas Basin sedimentary subsurface. Rapidly decreasing downcore cell densities create special challenges for DNA extraction due to declining biomass with depth.

Here we evaluate DNA and RNA extraction methods that proved workable for these deep subsurface sediments. We report recovery of DNA (Tables 1, 2) and RNA (Table 3) obtained by these protocols, and we discuss downcore trends in microbial cell density in Guaymas Basin sediment cores (post-cruise on-shore counts, Table 2) within the context of extraction yields. We provide qualified depth estimates when, based on current technical limitations, the recovery of DNA and RNA becomes insufficient to support microbial community analyses by PCR, and by metagenomic and metatranscriptomic sequencing. For DNA, we term these limits the “DNA event horizon” and suggest strategies to gradually extend them. For the Guaymas sample set, DNA recovery extends deeper into subsurface sediments than RNA recovery, which required additional washing steps to remove inhibitors. Due to the different extraction procedures for DNA vs. RNA and apparent sensitivity limits, the ranges of DNA and RNA recovery in the deep subsurface are not the same; therefore, we cannot identify a combined “DNA/RNA event horizon” at this time. Yet, this RNA extraction procedure should provide a basis for further development.

**Table 1.**
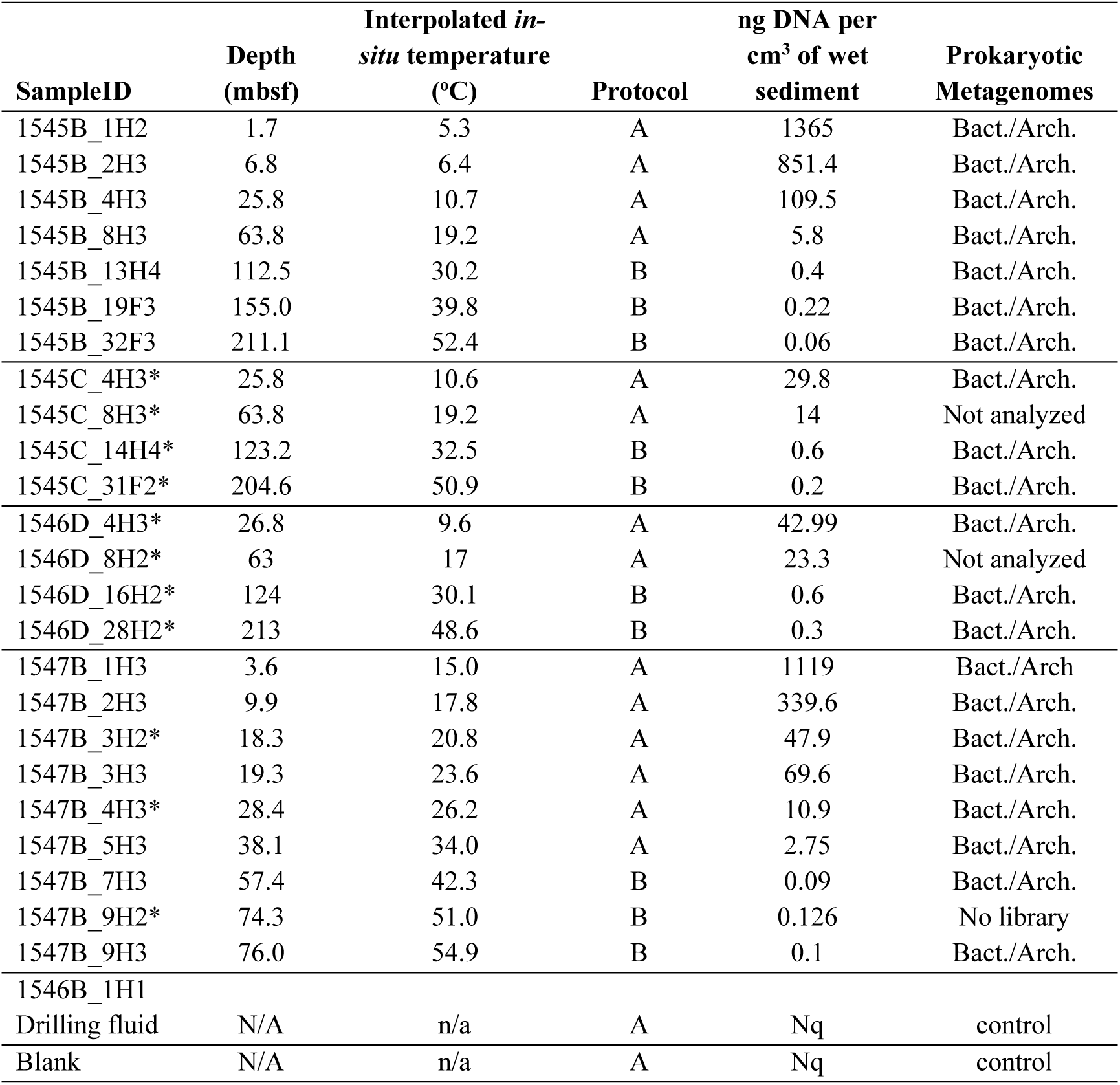
DNA yields from Guaymas subsurface sediments in sites U1545B and U1547B, extracted by G. Ramírez (no asterisk) and D. Bojanova (asterisk). The column “Prokaryotic Metagenomes” indicates samples used for metagenomic analyses (Bojanova et al., 2023, Mara et al., 2023b). The sequencing data obtained from our two controls (a drilling fluid sample and a kit control) were excluded from the metagenome analyses. n/a: not applicable; Nq.: DNA was not quantifiable using double strand high sensitivity DNA assays (Qubit); No library: unsuccessful library prep.

**Table 2.** DNA yields from Guaymas subsurface sediments, extracted for PCR assays by D. Beaudoin and P. Mara (Edgcomb Lab, WHOI) and J.E. Hinkle (U1548C samples; Teske Lab, UNC). Sample depths are noted as meters below seafloor (mbsf). An X in the general 16S and Archaeal 16S rRNA gene PCR column indicates that PCR amplification was successful. Nq: DNA was not quantifiable using double strand high sensitivity DNA assays (Qubit); da: PCR amplification was attempted but remained unsuccessful; ---: no PCR was performed. The *mcrA* gene PCR column is coded as follows: X indicates positive only for mcr-ANME1 primers; X1 indicates positive only for mcr-IRD primers targeting general methanogens but not ANME1, X2 indicates positive for both mcr-IRD and mcr-ANME1 primers. Cell counts were compiled for these samples and plotted in Figure 3.

| Sample ID | Depth<br>(mbsf) | Interpolated<br>*in-situ*<br>temperature<br>(°C) | ng DNA<br>per gr of<br>sediment | Sediment<br>extracted<br>(gr) | 400 bp<br>bacterial/arch<br>aeal 16S<br>rRNA gene<br>PCR | 800 bp<br>archaeal<br>16S rRNA<br>gene PCR | *mcrA*<br>gene<br>PCR | Cells<br>per ml | Potential<br>DNA yield,<br>1 fg DNA/cell<br>(ng ml<sup>-1</sup>) |
| --- | --- | --- | --- | --- | --- | --- | --- | --- | --- |
| 1545B_1H2 | 1.7 | 5.3 | 300.0 | 0.5 | X | X | da | 4.81x10<sup>8</sup> | 481 |
| 1545B_4H2 | 25.8 | 10.7 | 185.0 | 0.5 | X | X | X | 2.47x10<sup>7</sup> | 24.7 |
| 1545B_6H2 | 43.6 | 14.7 | 15.8 | 9.5 | X | X | da | 4.29x10<sup>6</sup> | 4.29 |
| 1545B_9H2 | 72.2 | 21.1 | 0.833 | 30 | X | --- | --- |  |  |
| 1545B_11H3 | 92.4 | 25.6 | 3.125 | 8 | X | X | da | 4.73x10<sup>5</sup> | 0.473 |
| 1545B_13H3 | 111.2 | 29.9 | 3.66 | 5 | X | X | da | 8.41x10<sup>5</sup> | 0.841 |
| 1545B_15H3 | 130.7 | 34.3 | 3.06 | 5 | X | X | da | 2.13x10<sup>5</sup> | 0.213 |
| 1545B_20F4 | 160.1 | 40.9 | 2.71 | 5 | X | X | da |  |  |
| 1545B_25F2 | 177.4 | 44.8 | 1.75 | 5 | da | X | da | 1.85x10<sup>4</sup> | 0.0185 |
| 1545B_30F2 | 200.7 | 50.1 | Nq | 5 | da | --- | --- |  |  |
| 1545B_34F3 | 219.5 | 54.3 | Nq | 5 | da | --- | --- |  |  |
| 1545B_44F2 | 266.7 | 64.9 | 1.94 | 5 | da | --- | --- |  |  |
| 1545B_50F3 | 289.6 | 70 | 0.69 | 5 | da | --- | --- |  |  |
| 1545B_58F2 | 315.8 | 75.9 | Nq | 5 | da | --- | --- |  |  |
| 1545B_60F2 | 325.1 | 78 | Nq | 5 | X | --- | --- |  |  |
| 1545B_64X2 | 359.3 | 85.7 | Nq | 5 | da | --- | --- |  |  |
| 1545A_62X2 | 393 | 93.3 | Nq | 5 | da | --- | --- |  |  |
| 1545A_72X2 | 485.5 | 114.1 | Nq | 5 | da | --- | --- |  |  |
| 1545A_74X3 | 500.2 | 117.4 | Nq | 5 | da | --- | --- |  |  |
| 1546B_1H2 | 0.8 | 2.8 | 900.0 | 0.5 | X | X | da | 1.30x10<sup>9</sup> | 1300 |
| 1546B_3H2 | 16.1 | 6.2 | 100 | 0.5 | X | X | da | 2.28x10<sup>8</sup> | 228 |
| 1546B_5H2 | 35.1 | 10.4 | 64.5 | 0.5 | X | X | da | 5.53x10<sup>6</sup> | 5.53 |
| 1546B_7H2 | 54.0 | 14.6 | 36.2 | 0.5 | X | X | da | 6.10x10<sup>6</sup> | 6.10 |
| 1546B_9H2 | 73.1 | 18.8 | 16.7 | 0.5 | X | X | da | 8.56x10<sup>5</sup> | 0.856 |
| 1546B_12H2 | 102.1 | 25.2 | 2.29 | 5 | X | X | da | 2.11x10<sup>5</sup> | 0.211 |
| 1546B_15H3 | 131.9 | 31.8 | 1.69 | 5 | da | X | da | 7.77x10<sup>4</sup> | 0.0777 |
| 1546B_20H3 | 168.8 | 40.0 | 1.26 | 5 | da | X | da | 8.68x10<sup>3</sup> | 0.00868 |
| 1546B_22H3 | 188 | 44.2 | 1.49 | 5 | da | --- | --- |  |  |
| 1546B_28F2 | 211.6 | 49.5 | 0.5 | 5 | da | --- | --- |  |  |
| 1546B_35F3 | 237.9 | 55.3 | 2 | 5 | da | --- | --- |  |  |
| 1546B_39F2 | 252.5 | 58.5 | 2.5 | 5 | da | --- | --- |  |  |
| 1546B_49F3 | 273.7 | 63.2 | Nq | 5 | da | --- | --- |  |  |
| 1546C_24R3 | 450.8 | 103.7 | Nq | 5 | da | --- | --- |  |  |
| 1546C_28R2 | 471.3 | 108.1 | Nq | 5 | da | --- | --- |  |  |
| 1546C_34R2 | 500.0 | 114.5 | Nq | 5 | da | --- | --- |  |  |
| 1546C_40R2 | 529.6 | 121.0 | Nq | 5 | X | --- | --- |  |  |
| 1547B_1H2 | 2.2 | 14.2 | 300 | 0.5 | X | X | da |  |  |
| 1547B_2H2 | 8.7 | 17.5 | 250 | 0.5 | X | X | X1 | 1.25x10<sup>8</sup> | 125 |
| 1547B_3H2 | 17.7 | 22.0 | 42.86 | 3.5 | X | X | da | 9.32x10<sup>6</sup> | 9.32 |
| 1547B_5H2 | 36.9 | 31.9 | 20.0 | 0.5 | X | X | da | 1.35x10<sup>6</sup> | 1.35 |
| 1547B_7H2 | 55.9 | 41.6 | 2.0 | 5 | da | X | da | 3.20x10<sup>5</sup> | 0.32 |
| 1547B_8H2 | 65.7 | 46.6 | 2.0 | 5 | da | X | X | 4.30x10<sup>4</sup> | 0.043 |
| 1547B_9H2 | 74.3 | 51.0 | 0.91 | 5 | da | X | X | 3.85x10<sup>4</sup> | 0.0385 |
| 1547B_12F2 | 94.6 | 60.7 | Nq | 5 | da | X | da | 1.22x10<sup>3</sup> | 0.0012 |
| 1547B_25F2 | 132.1 | 80.6 | Nq | 5 | da | X | da | 2.77x10<sup>3</sup> | 0.0028 |
| 1547B_28F2 | 146.2 | 87.8 | Nq | 5 | da | --- | --- |  |  |
| 1547B_29F2 | 151.0 | 90.2 | Nq | 5 | da | --- | --- |  |  |
| 1548B_1H2 | 2.1 | 8.2 | 700 | 0.5 | X | X | X | 5.29x10<sup>8</sup> | 529 |
| 1548B_2H3 | 8.9 | 13.7 | 135 | 0.5 | X | X | X | 9.96x10<sup>7</sup> | 99.6 |
| 1548B_3H4 | 20.4 | 22.9 | 33.9 | 0.5 | X | X | X2 | 7.68x10<sup>7</sup> | 76.8 |
| 1548B_4H7 | 33.5 | 33.5 | 10 | 0.5 | X | X | da | 1.78x10<sup>7</sup> | 17.8 |
| 1548B_5H5 | 39.6 | 38.3 | 10 | 0.5 | X | X | da | 7.52x10<sup>6</sup> | 7.52 |
| 1548B_6H2 | 46.2 | 43.6 | 5.9 | 0.5 | X | X | da | 1.17x10<sup>6</sup> | 1.17 |
| 1548B_7H3 | 56.4 | 51.8 | nq | 5 | X | --- | --- |  |  |
| 1548B_8H5 | 69.5 | 62.4 | 1 | 5 | da | X | da | 1.01x10<sup>3</sup> | 0.001 |
| 1548B_9H3 | 76.5 | 68.0 | 2.5 | 5 | X | da | X | 1.62x10<sup>3</sup> | 0.0016 |
| 1548C_1H3 | 3.4 | 19.2 | 256 | 0.5 | --- | --- | --- |  |  |
| 1548C_2H3 | 10.6 | 26.1 | 146 | 0.5 | X | --- | --- |  |  |
| 1548C_3H6 | 24.2 | 39.1 | 21.2 | 0.5 | X | --- | --- |  |  |
| 1548C_5H2 | 37.5 | 51.9 | 1.24 | 5 | X | --- | --- |  |  |
| 1548C_5H4 | 40.4 | 54.6 | 3.8 | 0.5 | --- | --- | --- |  |  |
| 1548C_6H4 | 50.5 | 64.3 | 0.71 | 5 | X | --- | --- |  |  |
| 1549B_1H2 | 1.6 | 3.5 | 1040 | 0.5 | X | X | da | 7.25x10<sup>8</sup> | 725 |
| 1549B_2H2 | 7.0 | 4.6 | 364 | 0.5 | --- | X | da | 4.92x10<sup>8</sup> | 492 |
| 1549B_3H2 | 16.5 | 6.4 | 161 | 0.5 | X | X | da |  |  |
| 1549B_6H3 | 45.6 | 12.1 | 63.5 | 0.5 | X | X | X | 2.48x10<sup>7</sup> | 24.8 |
| 1549B_9H3 | 74.4 | 17.6 | 23.8 | 5 | X | X | da |  |  |
| 1549B_12H3 | 103.7 | 23.3 | 0.73 | 5 | X | X | da | 3.69x10<sup>6</sup> | 3.69 |
| 1549B_15H4 | 133.4 | 29.1 | 5 | 5 | X | X | da | 7.73x10<sup>5</sup> | 0.773 |
| 1549B_18H4 | 161.4 | 34.5 | 3.25 | 5 | X | --- | --- |  |  |
| 1550B_1H2 | 2.0 | 3.8 | 1150 | 0.5 | X | X | X | 2.85x10<sup>8</sup> | 285 |
| 1550B_3H2 | 16.9 | 5.8 | 465 | 0.5 | X | X | X | 5.71x10<sup>7</sup> | 57.1 |
| 1550B_7H2 | 54.8 | 10.9 | 96 | 0.5 | X | X | X2 | 1.82x10<sup>7</sup> | 18.2 |
| 1550B_11H2 | 92.3 | 16.0 | 4.75 | 5 | da | da | da |  |  |
| 1550B_19X2 | 142.0 | 22.7 | nq | 5 | da | da | X | 1.04x10<sup>2</sup> | 0.0001 |
| 1551B_1H1 | 0.8 | 4.8 | 700 | 0.5 | X | X | X | 4.18x10<sup>8</sup> | 418 |
| 1551B_2H2 | 5.8 | 5.3 | 454 | 0.5 | --- | X | X | 2.97x10<sup>8</sup> | 297 |
| 1551B_3H2 | 15.4 | 6.3 | 103 | 0.5 | X | X | X | 1.38x10<sup>7</sup> | 13.8 |
| 1551B_5H2 | 34.2 | 8.2 | 30 | 0.5 | X | X | X | 3.26x10<sup>6</sup> | 3.26 |
| 1552B_1H2 | 0.8 | 3.8 | 1100 | 0.5 | X | X | X | 2.99x10<sup>8</sup> | 299 |
| 1552B_3H3 | 19.2 | 8.6 | 605 | 0.5 | --- | X | X | 8.57x10<sup>6</sup> | 8.57 |
| 1552B_3H4 | 20.4 | 8.9 | 129 | 5 | X | da | da |  |  |
| 1552B_6H2 | 46.9 | 15.8 | 34.4 | 0.5 | X | X | da | 2.49x10<sup>6</sup> | 2.49 |

**Table 3.** RNA recovery from samples that yielded metatrancriptomes (Mara et al. 2023a). Nq: no quantifiable RNA using Qubit RNA high sensitivity assays (Qubit).

| Site | SampleID | Depth (mbsf) | Interpolated *in-situ* temperature (°C) | ng RNA per gr of sediment | Sediment extracted (gr) |
| --- | --- | --- | --- | --- | --- |
| U1545B | U1545B-1H2 | 1.7 | 5.3 | 38.2 | 11 |
|  | U1545B-4H3 | 25.8 | 10.7 | 12.5 | 12 |
|  | U1545B-11H3 | 92.4 | 25.6 | Nq | 11 |
| U1546B | U1546B-1H2 | 0.8 | 2.8 | 63.8 | 11 |
|  | U1546B-3H2 | 16.1 | 6.2 | 10.5 | 12 |
|  | U1546B-12H2 | 102.1 | 25.2 | Nq | 13 |
| U1547B | U1547B-1H2 | 2.2 | 14.2 | 22.2 | 13.5 |
|  | U1547B-5H2 | 36.9 | 31.9 | 9 | 10 |
|  | U1547B-9H2 | 74.3 | 51 | Nq | 12 |
| U1548B | U1548B-1H2 | 2.1 | 8.2 | 27.2 | 13 |
|  | U1548B-4H7 | 33.5 | 33.5 | nq | 14 |
| U1549B | U1549B-1H2 | 1.6 | 3.5 | 42.5 | 13 |
|  | U1549B-3H2 | 16.5 | 6.4 | Nq | 10 |
| U1550B | U1550B-1H2 | 2.0 | 3.8 | 42 | 10 |
|  | U1550B-3H2 | 16.9 | 5.8 | 14.1 | 13 |
| U1551B | U1551B-1H1 | 0.8 | 4.8 | 16.5 | 13.5 |
|  | U1551B-3H2 | 15.4 | 6.3 | Nq | 11 |
| U1552B | U1552B-1H2 | 0.8 | 3.8 | 48.8 | 12 |
|  | U1552B-3H4 | 20.4 | 8.9 | 16.8 | 12 |

## Materials and Methods

### DNA extraction

DNA was extracted from selected core samples using FastDNA™ SPIN Kit for Soil (MP Biomedicals). For each sample listed in Table 1, three parallel (triplicate) DNA extractions were performed. The three extracts were pooled together and concentrated as discussed below. The samples listed in Table 2 were extracted once (single extraction). Otherwise, sediments were processed following the manufacturer’s protocol with homogenization modifications as described previously (Ramírez et al., 2018). We note that some DNA extraction sets (Table 1, Figure 1) started with volumetrically defined wet sediment samples of 0.5 cm^3^ each, whereas others started with weighed wet sediment samples of 0.5 g each (Table 2, Figure 2). These concentrations can be converted into each other to an approximate degree by using a factor of 1.7 g/cm^3^, the average wet bulk density of an extensive set of IODP sediments (Tenzer and Gladkikh 2014). The extraction procedure tolerates a sediment input of 0.5 cm^3^, which weighs more than 0.5 g. Procedures should be kept internally consistent for each sample set, and the extraction vials have to be balanced during the centrifugation steps.

**Figure 1.**
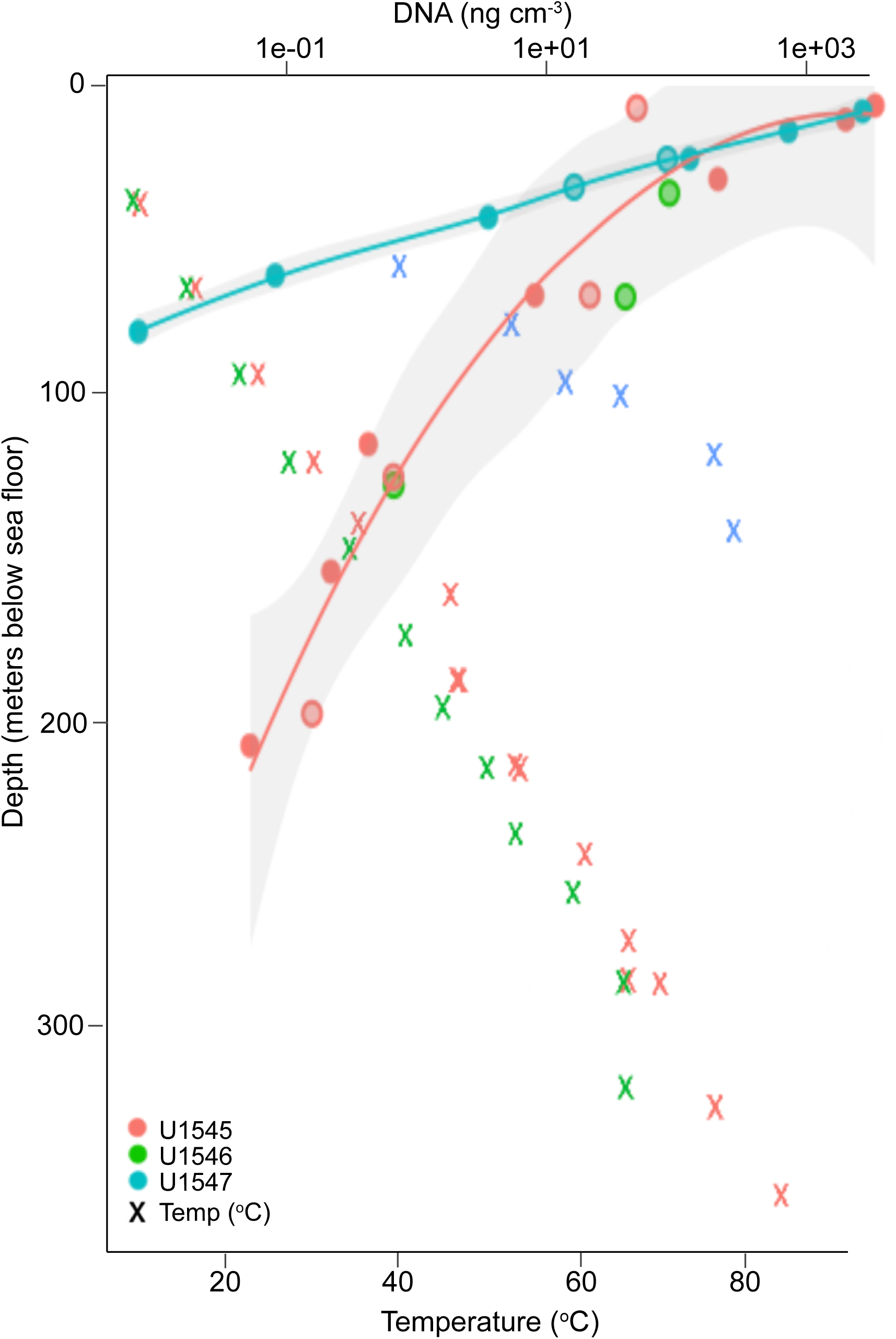
Downcore DNA concentration profiles for Holes U1545B and U1547B, extracted for metagenomic sequencing of bacteria and archaea. The temperature values represent the in-situ measurements (Teske et al. 2021b, d). The plotted lines for DNA concentrations follow a generalized additive model of exponential functions, flanked by 95% confidence intervals (ggplot2 smooth_stat() function) that are shown as grey zones. Extractions give consistent results across different sample sets, as shown by solid circles representing initial DNA extractions by G. Ramirez, and hollow circles representing a second set of DNA extractions by D. Bojanova.

**Figure 2.**
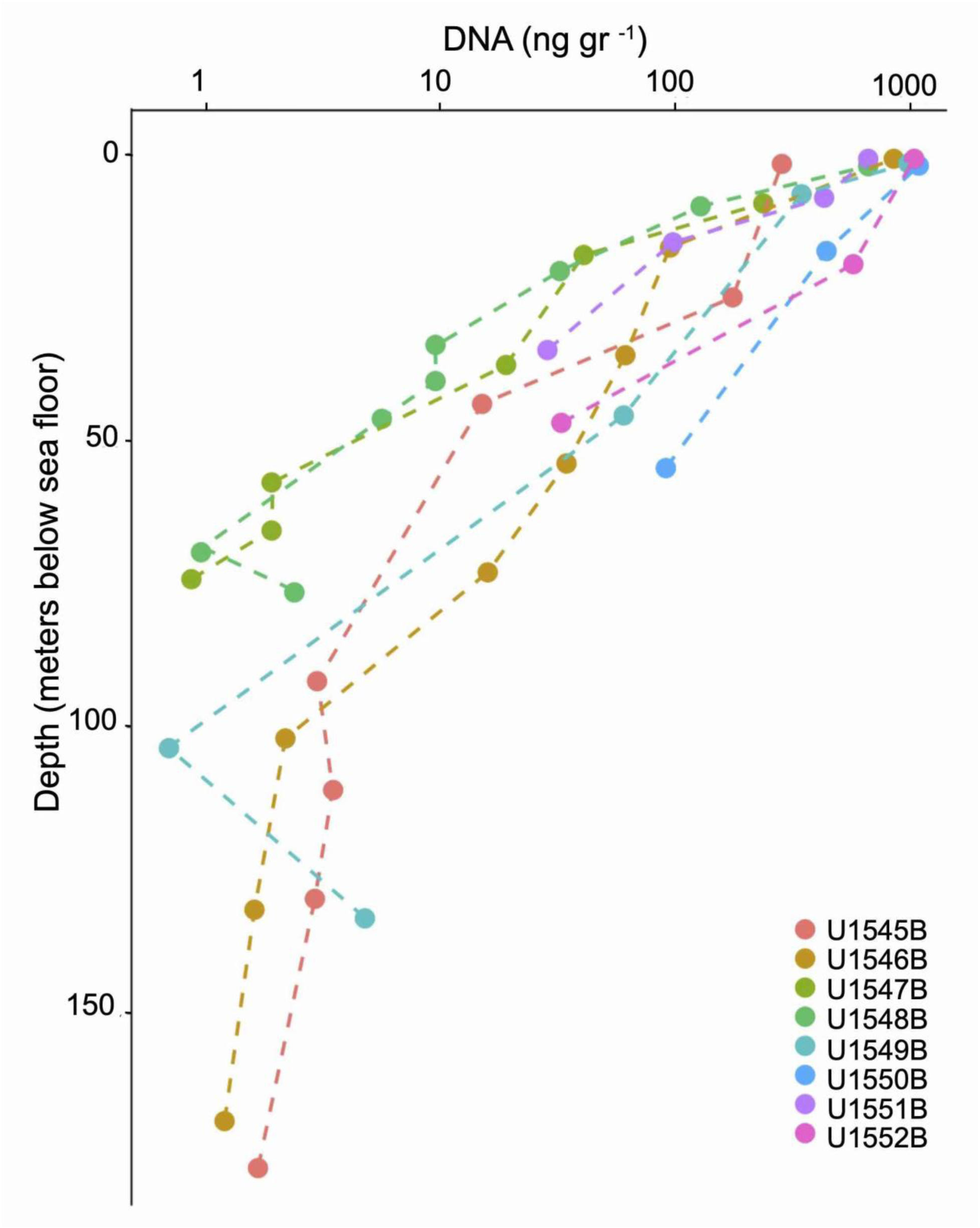
Downcore DNA concentration profiles for Holes U1545B to U1552B, extracted for PCR amplification of 16S rRNA genes and functional genes (Table 2).

Briefly, each sediment sample was homogenized twice (vs. the manufacturers’ suggestion of a single homogenization step) in Lysing Matrix E tubes for 40 seconds at speed 5.5 m/s, using the MP biomedicals bench top homogenizer equipped with 2 ml tube adaptors. Between the two homogenization rounds the samples were placed on ice for 2 minutes. After the second homogenization the samples were centrifuged at 14,000 x g for 5 minutes following the manufacturer’s suggestion. In the next step, which was a second modification from the standard protocol (applied only to DNA extractions of Table 1), the supernatant and the top layer of the pellet was transferred to a clean 2 ml tube where proteins were precipitated by the addition of the protein precipitation solution (PPS) provided in the extraction kit. The rest of the extraction protocol followed the manufacturer’s recommendations. When parallel extractions were performed, the extracts were pooled and concentrated using EMD 3kDa Amicon Ultra-0.5 ml Centrifugal Filters (Millipore Sigma). The DNA concentration was fluorometrically measured using the High Sensitivity (HS) double strand (ds) DNA Qubit assays (Qubit™ dsDNA HS and BR Assay Kits). A kit control extraction (extraction with no sediment added; blank or “witness” extraction) was included to account for any potential kit and handling contaminants. For extractions from deeper sediments where triplicate sample pooling did not result in quantifiable amounts of DNA, we performed 10 parallel extractions which were subsequently pooled and concentrated using the Amicon filters as described above.

**<u>Protocol A</u>** is used for high biomass samples with expected cell densities ≥ 10^6^ cells per cm^3^. Three parallel extractions are done per sample, including the blank or “witness” extractions.

1. Add up to 0.5 cm^3^ sediment or 0.5 g sediment to a Lysing Matrix E tube.
2. Add 978 μl Sodium Phosphate Buffer to sample in Lysing Matrix E tube.
3. Add 122 μl of MT buffer.
4. Homogenize in the FastPrep homogenizer or equivalent beat-beating homogenizer for 40 seconds twice, at speed setting of 5.5 m/s with 2-minute rest periods on ice.
5. Centrifuge at 14,000 x g for 5 mins to pellet debris.
6. Transfer supernatant to a clean 2 ml tube. *[Note: At this step, a modification of the manufacturer’s protocol is recommended. Carefully aspirate a few (50-100 μl max) of the top part of the pellet as well. Avoiding the pellet leads to significantly lower DNA yields, meaning that the positive charges on pelleted particles sequester DNA. After bead-beating, DNA is extracellular and fully exposes its negatively charged phosphodiester backbone. This trapped DNA is subsequently released into Sodium Phosphate Buffer.]*
7. Add 250 μl PPS (protein precipitating solution) and mix by shaking by hand 10 times.
8. Centrifuge at 14,000 x g for 15 minutes to precipitate the pellet. *[Note: Due to the intentional pellet carry over from step 6, 15 minutes are allowed for protein precipitation]*. Transfer the supernatant into a clean 15 ml Falcon tube.
9. Resuspend the Binding Matrix suspension and add 1 ml to the supernatant in the 15 ml Falcon tube.
10. Place on rotator or invert by hand for 2 minutes to allow binding of DNA. Place tube in rack for 3 minutes to allow settling of the Binding Matrix.
11. Remove and discard 500 μl of supernatant being careful to avoid settled Binding Matrix.
12. Resuspend Binding Matrix in the remaining amount of supernatant. Transfer approximately 600 μl of the mixture to a SPIN Filter and centrifuge at 14,000 x g for 1 minute. Empty the catch tube, add 600 μl of the mixture and centrifuge again at 14,000 x g for 1 minute. Repeat process until all mixture passes through the SPIN filter.
13. Add 500 μl prepared SEWS-M (Nucleic Acid Wash Solution provided by the kit) and gently resuspend the Binding Matrix pellet using the force of the liquid from the pipet tip. Ensure that absolute ethanol has been added to the concentrated SEWS-M stock solution as suggested by the manufacturer.
14. Centrifuge at 14,000 x g for 1 minute. Empty the catch tube.
15. Centrifuge again at 14,000 x g for 2 minutes to “dry” the Binding Matrix from residual wash solution. Discard the catch tube and replace with a new, clean catch tube.
16. Air dry the SPIN Filter for 5 minutes at room temperature.
17. Gently resuspend the Binding Matrix (collected inside the SPIN filter) in 50 μl of DES (DNase/Pyrogen-free water). [*Note: not 30 μl of DES, in manufacturer’s protocol*].
18. Centrifuge at 14,000 x g for 1 min to elute DNA from the Binding Matrix into the clean catch tube. Discard the SPIN filter. This will result in ∼50 μl of DNA in the catch tube. *[NOTE: Use the smallest possible volume (1 μl) for Qubit fluorometric quantification to save DNA]*. Store all extracts, or single pooled extracts at −80°C.

**<u>Protocol B</u>** is used for low biomass samples with expected cell densities are < 10^6^ cells per cm^3^. 10 parallel extractions were performed per sample and include a batch of blank or “witness” extractions.

Steps 1 to 18 are the same as in Protocol A, performed 10 times in parallel.

19. Potential stopping point or break: Store all extracts, or single pooled extracts at −80°C now if necessary.
20. Assuming that yields are not sufficient (which is nearly always the case if cell concentrations are expected < 10^6^ cells per cm^3^ in the sample) transfer the full combined volume of all 10 individual extractions (∼500 μl) to an Amicon Ultra 0.5 30K filter for concentrating the DNA, following the manufacturer’s instructions.

**<u>Protocol C</u>** improves the DNA yield by adding a Lysozyme and Proteinase K step before performing DNA with FastDNA^TM^ SPIN kit for soil (MP Biomedical). Protocol C was tested with a shallow subsurface sediment from 0.8 mbsf (site U1546B), and a method control. One 0.5-gram sediment sample was pretreated with Lysozyme and Proteinase K, and the other sample was not. The sample with the lysozyme and the proteinase K pretreatment before performing extraction Protocol A, gave higher DNA yield when compared to the non-pretreated sediment (0.8 ng µl^-1^ vs. 0.2 ng µl^-1^ fluorometrically quantified using Qubit™ dsDNA HS). The method control did not yield DNA. The applied heating steps (37°C and 55°C for 45 and 20 minutes, respectively) improved the DNA extraction efficiency but also seemed to remove inhibitory compounds that interfere with DNA extraction (e.g., humic acids, hydrocarbons). The protocol could be scaled up and tested further in triplicate or tenfold extractions as described in Protocol B.

1. Prepare 50 mM lysozyme solution by diluting 2 mg of lysozyme in 40 μl of Sodium Phosphate Buffer provided by the FastDNA^TM^ SPIN kit for soil (MP Biomedical).
2. Prepare 10 mM proteinase K solution by diluting 1 mg of Proteinase K in 100 μl of Sodium Phosphate Buffer provided by the FastDNA^TM^ SPIN kit for soil (MP Biomedical).
3. Add up to 500 mg of sediment sample to a Lysing Matrix E tube.
4. Add 800 μl Sodium Phosphate Buffer to sample in Lysing Matrix E tube and the 40 μl of the lysozyme solution.
5. Incubate at 37°C rotating for 45 minutes. Incubation time can increase up to 1 hr.
6. After lysozyme treatment add 100 μl proteinase K solution and 100 μl of 20% SDS (w/v) to the sample in the Lysing Matrix E tube. The addition of 20% SDS can be replaced by 120 μl MT buffer provided by the kit. [*Note: SDS concentration in MT buffer is undisclosed, but SDS is known to stimulate the activity of proteinase Keven at low concentrations between 0.5-2% w/v (Hilz et al., 1975).*] Replacement of 20% SDS with 120 μl MT was also tested and worked.
7. Incubate at 55°C rotating for 20 minutes. Incubation time can increase up to 40 minutes.
8. Place the Lysing Matrix E tube with the sample in the FastPrep homogenizer for 40 seconds at speed setting of 6 m/s.
9. Continue with the DNA extraction protocol A starting from step 5 until step 18.
10. Assuming that DNA yields are not sufficient, then continue with steps 19 to 20 described in Protocol B.

### Other methods

We also tested DNA isolation with the Qiagen DNAeasy Powersoil Pro kit, but obtained an order of magnitude lower DNA yields compared to the FastDNA kit. Three samples (U1545C_4H_3, U1545C_8H_3, and U1547B_3H_3) yielded 2.876, 0.684 and 4.648 ng DNA/g sediment with the Qiagen kit, and 164.25, 8.66 and 104.4 ng/g sediment with the FastDNA kit, respectively. Likewise, DNA extractions using higher sediment volumes (up to 10 grams) with Qiagen DNeasy PowerMax Soil led also to low DNA yields (similar ranges as those reported for Qiagen DNAeasy Powersoil Pro kit tests). Because higher volumes of sediment also increase the concentrations of inhibitory compounds found in Guaymas subsurface sediments, our teams concluded that small-scale (0.5 cm^3^ or 0.5 g) sediment extractions offer a more favorable balance of DNA yield and inhibitor accumulation. These results confirmed the decision of all parties to use the modified FastDNA procedure as outlined above in protocols A or B.

### RNA extraction

To demonstrate microbial activity and associated gene expression, RNA extraction, sequencing and metatranscriptomic analyses are essential. Prior protocols for deep subsurface total RNA extraction from our labs involved bead-beating, organic extraction and ethanol precipitation (Sørensen and Teske 2006; Biddle et al. 2006) or used soil RNA extraction kits that excel also for samples that have high humic content, including sediments (Edgcomb et al. 2011, Orsi et al., 2013a,b). However, for over a year of effort, Guaymas Basin subsurface sediments maintained at −80°C defeated all attempts at RNA extraction using various methods, including also the original RNA isolation protocols published by Chomczynski and Sacchi (1987). Aside from the different RNA isolation protocols, we also performed different sediment pretreatments that have been shown to increase cell extraction efficiency from deep subsurface sediments (Kallmeyer et al., 2008). Yet, even with those pretreatments (e.g., acidification of sediments for 10 minutes with 0.43 M sodium acetate to dissolve carbonates), RNA could not be isolated. The steps that eventually led to successful RNA extraction involved initially washing sediment samples twice with absolute ethanol (200 proof; purity ≥ 99.5%; Thermo Scientific Chemicals), followed by one wash with DEPC-treated water (Fisher BioReagents) before extracting RNA. This procedure appears to remove sufficient hydrocarbons and other inhibitory elements present in Guaymas sediments. Similar washing procedures have been tested for other hydrocarbon-rich sediments (Lappé and Kallmeyer 2011). Without ethanol and DEPC water washes, all attempts resulted in low or zero RNA yield. At the time of writing this manuscript, all metatranscriptomic analyses of Guaymas Basin subsurface sediments (Mara et al., 2023a, Mara et al., 2023b) depended on including these washing procedures. We also note that washing steps might be applied to future DNA extraction protocols, to remove inhibitors that accumulate when extraction volumes are scaled up.

In brief, 10-15 grams of frozen Guaymas Basin sediments were transferred into UV-sterilized 50 ml Falcon tubes (RNAase/DNase free) using clean, autoclaved and ethanol-washed metallic spatulas. Each tube received an equal volume of absolute ethanol and was shaken manually for 2 minutes followed by 30 seconds of vortexing at full speed to create a slurry. Samples were transferred into an Eppendorf centrifuge (5810R) and were centrifuged at room temperature for 2 minutes at 2000 rpm. The supernatant was decanted, and the ethanol wash was repeated. After decanting the supernatant of the second ethanol wash, an equal volume of DEPC water was added into each sample. Samples were manually shaken and vortexed as before to create slurry and were transferred into the Eppendorf centrifuge (5810R) where they were centrifuged at room temperature for 2 minutes at 2000 rpm. The supernatant was decanted, and each sediment sample was immediately divided into three bead-containing 15 mL Falcon tubes, provided by the PowerSoil Total RNA Isolation Kit (Qiagen). RNA was extracted as suggested by the manufacturer with the modification that the RNA extracted from the three aliquots was pooled into one RNA collection column and eluted at 30 μl final volume. All RNA extractions were performed in a UV-sterilized clean hood (two UV cycles of 15 min each) that was installed with HEPA filters. Surfaces inside the hood and pipettes were sterilized with RNase AWAY (Thermo Scientific) before every RNA extraction and in between extraction steps. Trace DNA contaminants were removed from RNA extracts using TURBO DNase (Thermo Fisher Scientific) and the manufacturer’s protocol. Carryover DNA removal from the RNA extracts was confirmed with PCR reactions using primers for the small ribosomal subunit of 16S rRNA gene [BACT1369F: 5’CGGTGAATACGTTCYCGG3’ and PROK1541R: 5’AAGGAGGTGATCCRGCCGCA 3’; Suzuki et al., 2000)]. Each 25 μl PCR reaction was prepared using GoTaq G2 Flexi DNA Polymerase (Promega) and contained 0.5 U μl^−1^ GoTaq G2 Flexi DNA Polymerase, 1X Colorless GoTaq Flexi Buffer, 2.5 mM MgCl_2_, (Promega) 0.4 mM dNTP Mix (Promega), 4 μM of each primer (final concentrations), and DEPC water. Further, to confirm absence of DNA contamination due to handling and PCR reagents, all PCR experiments included negative controls (blanks) where no DNA was added. PCR amplifications were performed in an Eppendorf Mastercycler Pro S Vapoprotect (Model 6321) thermocycler with following conditions: 94°C for 5 min, followed by 35 cycles of 94°C (30 s), 55°C (30 s), and 72°C (45 s). The PCR reaction products were run in 2% agarose gels (Low-EEO/Multi-Purpose/Molecular Biology Grade Fisher BioReagents) to confirm absence of DNA products. RNA quantification (ng μl^-1^) was performed using Qubit RNA High Sensitivity (HS), Broad Range (BR), and Extended Range (XR) Assay Kits, (Invitrogen). Because of the essential sediment washing steps with ethanol and DEPC-treated water, and the small volume of final RNA extractions (30 μL), the RNA integrity (RIN) of the extracted RNA was not estimated. The washes with absolute ethanol can enhance the already high susceptibility of RNA to degradation at room temperature, and therefore we selected to preserve the small volume of our RNA extracts for other mandatory steps involved in cDNA library preparation. These steps included quantifying the total RNA concentration before the cDNA library preparation, performing PCR reactions to confirm the absence of carryover DNA in the RNA extracts, and maintaining the necessary initial RNA volume (up to 10 μl) which is required as template for the synthesis of the single/first cDNA strand.

Amplified cDNAs from the DNA-free RNA extracts were prepared using the Ovation RNA-Seq System V2 (Tecan) following the manufacturer’s suggestions. All steps from RNA extraction through cDNA preparation were completed on the same day to avoid freeze/thaw cycles that might damage the integrity of RNA strands. cDNAs were submitted to the Georgia Genomics and Bioinformatics Core facility for library preparation and sequencing using NextSeq 500 PE 150 High Output (Illumina). Sequencing of the cDNA library prepared from the control sample (laboratory reagent control) was unsuccessful as it failed to generate any sequences that met the minimum length criterion of 300-400 base pairs.

While this RNA extraction and transcription protocol ran into downcore detection limits (Table 3), we note that every living subsurface cell with intact genomic DNA requires at least a minimum level of gene expression in order to survive and function, and thus we regard RNA extraction and detection limits foremost as methodological issues, not as evidence of fundamental constraints on life.

## Results

### Downcore trends in DNA yield

To obtain DNA for metagenomic analyses, protocols A and B were applied to DNA extractions from Site U1545B, U1545C, U1546D, and U1547B sediment samples, starting with 0.5 cm^3^ wet sediment samples. DNA yields are tabulated in Table 1 and plotted in Figure 1. The yield of DNA extractions decreases exponentially with sediment depth and temperature. In near-surface sediments, DNA yields are high, around 1000 to 1500 ng DNA, or 1 to 1.5 µg DNA per cm^3^ wet sediment. DNA yields decrease towards the limit of reliable detection and quantification by Qubit (below 0.1 ng per cm^3^) within 75 mbsf for site U1547B at Ringvent, and within ca. 200 mbsf at Site U1545B in the Northwestern Guaymas Basin. The steep thermal gradient at Ringvent means that a depth of 75 mbsf corresponds to in-situ temperatures near 50°C to 55°C whereas the cooler sediments in northwestern Guaymas Basin reach these temperatures only near 200 mbsf (Table 1 and Figure 1). The temperature-related differences in DNA recovery impact the depth horizons for successful metagenomic sequencing, which generally requires nanogram amounts of DNA. For example, generating Illumina libraries from low-biomass samples requires a previously reported minimum of 3.65 ng DNA (Jiang et al. 2015).

Independently, Protocols A and B were applied to extract DNA for another sequencing project from all sites; here extractions started with 0.5 g wet weight samples (Table 2, Figure 2). The DNA concentrations show a similar decline over several orders of magnitude as observed for the previous set of extractions. In the northwestern sites U1545 and U1546, DNA concentrations decline over almost three orders of magnitude, from the µg to the ng range per g sediment, within the upper 170 to 190 mbsf (40-45°C). DNA yields decline over three orders of magnitude at the Ringvent sites U1547 and U1548, with this sharp decline occurring within the upper 70 to 80 mbsf (50-60°C); some uncertainty for the deeper samples is not resolved since DNA concentrations dip below the 0.1 ng limit of Qubit detection (Table 2). In the cold seep-affiliated site U1549 and the axial site U1550, DNA concentrations decline by three orders of magnitude within approx. 140 mbsf (25-30°C). The shorter DNA gradients of Southeastern site U1551 and Northern seep site U1552 drop by more than one or two orders of magnitude over depths of 35 and 45 meters, respectively, and remain entirely within cool temperatures not exceeding 16°C (Table 2).

### Determination of the “DNA event horizon”

Empirically, the current limit for contaminant-free DNA that is consistently suitable for metagenomic analysis of Guaymas Basin sediments is around 0.2 to 0.5 ng extracted DNA/cm^3^ wet sediment (Table 1). Lower DNA concentrations near 0.1 ng/cm^3^ may still yield metagenomes, but contaminants hypothesized to emerge from the reagent’s “kitome” (Salter et al. 2014) increasingly appear once DNA concentrations decrease to 0.1 ng extracted DNA/cm^3^. Assuming a cellular DNA content of approximately 1 to 2 femtogram (10^-15^g) DNA per cell (Bakken and Olsen, 1989; Button et al., 2001), 0.2 to 0.5 ng (10^-9^ g) DNA is the equivalent of 0.2 to 0.5 x 10^6^ cells. Therefore, given the complexity of our sample set, DNA of sufficient quantity and quality for metagenomic library construction and sequencing can be extracted from sediments with cell densities of ca. 0.2 to 0.5 x 10^6^ cells/cm^3^, more than 3 orders of magnitude below near-surface cell densities of 10^9^ cells/cm^3^ sediment. From the perspective of obtaining sufficient DNA for metagenomes, this is the current “DNA Event horizon” for this complex environment given our methodological approaches. Although this metagenomic DNA limit lags much behind the greater sensitivity of cell counts, it has improved considerably compared to earlier attempts to obtain subsurface metagenomes from non-amplified DNA extracts, which succeeded only for shallow subsurface sediments (Biddle et al. 2008, 2011). We also note that the successful preparation of metagenomic libraries from small amounts of DNA is crucial, requiring in some cases extra efforts at library preparation, and in this regard not all sequencing facilities may be equally capable. The participants of this study had their metagenomic libraries prepared at the DNA Sequencing and Genotyping Center of the University of Delaware.

The “DNA event horizon” is slightly different for targeted PCR (Polymerase Chain Reaction) amplification of specific genes, and depends not only on drilling site and depth but also on the amplification target (Table 2). PCR allows for the amplification of targeted gene sequences within dilute DNA extracts, increasing the sensitivity of detection. DNA extracts from all drilling sites were used for PCR amplification of bacterial & archaeal 16S rRNA gene segments with primer combinations of 515F-Y (Parada et al. 2015) and 926R (Quince et al. 2011), and partial archaeal 16S rRNA genes using primer combination 25F and 806R (Mara et al., 2023), and for *mcrA* genes (Hinkle et al., 2023) using primer combinations mcrIRD and mcrANME1 (Lever and Teske 2015). In contrast to the metagenomic extractions in Table 1, these DNA extractions for PCR were performed in single samples rather than in triplicate, and the sample size (0.5 g) was based on wet weight, not volume. Ten replicate extractions (Protocol B) were performed for a second PCR attempt if the initial PCR did not work (Table 2).

In our PCR survey, DNA concentrations near 1-2 ng DNA/g wet sediment were generally required for positive PCR outcomes (Table 2), whereas DNA samples at lower concentrations – which includes the previously defined metagenome sequencing limit (< 0.2 to 0.5 ng DNA/cm^3^, or below 0.34 to 0.85 ng DNA/g wet weight sediment, assuming a conversion factor of 1.7) – did not produce PCR amplicons of 16S rRNA genes or other genes. The only exceptions were low-DNA extracts from deep, hot Ringvent samples (U1547B) that yielded archaeal PCR products (Table 2). This surprising result – slightly higher DNA concentration requirements for PCR than for metagenomic sequencing – may reflect the fact that metagenomic library preparation does not select for specific genes, which necessarily account only for a very small proportion of the DNA pool. In contrast, PCR assays for 16S rRNA genes and functional genes pick out specific genes by design, like the proverbial needle in the haystack. Interestingly, PCR amplification of an 800 bp archaeal 16S rRNA gene segment was more consistently successful than amplification of shorter (300 bp) 16S rRNA gene segments using general prokaryotic (bacterial + archaeal) primers. Archaeal 800 bp amplicons were consistently recovered until depths of ca. 75 mbsf at Ringvent sites U1547B and U1548B, and 170 mbsf at the Northwestern sites U1545B and U1546B (Table 2). PCR amplification of mcrA genes with the ANME1-targeted primers gave positive results generally near and within methane-sulfate interfaces (Hinkle et al., 2023). Amplification of mcrA gene for methanogens worked only for few samples, consistent with the apparent rarity of methanogens in the Guaymas Basin subsurface (Bojanova et al., 2023). The robustness of Archaeal 16S rRNA gene-directed PCR assays over a wide spectrum of DNA concentrations and depths may result from several factors: the absence of Archaea from common laboratory and kit contaminants (Salter et al. 2014), the highly conserved Archaea-specific primer sites that contrast with the more ambiguous primer regions used for general prokaryotic (bacterial and archaeal) 16S rRNA gene surveys, and the downcore increasing relative proportion of archaea in the microbiome of hydrothermal sediments (Ramírez et al. 2021, Lagostina et al. 2021).

### Guaymas Basin and the Cragg line

In cold marine sediments from passive continental margins, such as the Peru Margin, microbial cell numbers decline very gradually, and successful DNA and RNA recovery and subsequent sequencing surveys extend to considerable depth (Inagaki et al., 2006; Biddle et al., 2006; Pachiadaki et al. 2016). To track cell abundance over three orders of magnitude, from 10^9^ cells per cm^3^ at the surface towards 10^6^ cells per cm^3^ at depth, required drilling and sampling down to 1000 meters sediment depth (Parkes et al., 2014). The gradual downcore decline in cell density for organic-rich, cold sediments follows a log-log regression line defined as Log cells = 8.05 – 0.68 Log depth (R^2^ = 0.70 and n = 2037; see Figure 2 in Parkes et al. 2014). We refer to this relationship as the Cragg line, in memory of cell count pioneer Barry Cragg. Interestingly, published cell counts from hydrothermal sediments (Cragg et al. 2000, Expedition 311 Scientists, 2011; Heuer et al. 2017) are significantly lower than their counterparts from cold sediments, and fall – regardless of considerable scatter – below the 95% confidence intervals of the Cragg line (Parkes et al., 2014). Similar trends appear in Guaymas Basin. Plotting Expedition 385 cell counts (with DNA stain SYBR Green I) shows that cells densities from all drilling sites and holes decrease by 3 orders of magnitude, from 10^9^ cells/cm^3^ sediment near the surface towards 10^6^ cells/cm^3^ sediment between ca. 50 and 100 m depth, and intersect with the Cragg line at ca. 50 m depth (Figure 3). This downcore decreasing trend is accelerated at the two hot Ringvent sites U1547 and U1548, and delayed at the cool sites 1549 and 1550 (Figure 3). Similarly, DNA yields in the Guaymas subsurface decline by three orders of magnitude by 50 to 100 m depth (Figure 1, Figure 2). In non-hydrothermal sediments such a decline occurs over well before 1000 m depth (Parkes et al. 2000). We see declining DNA yields already in sediments with warm in-situ temperatures of ca. 35-40°C (at ca. 40-50 m depth at Ringvent and 130-150 m depth at U1545), far below the temperature range for hyperthermophilic vent archaea (80°C and up), indicating that the survival of most subsurface microorganisms is already noticeably reduced at those depths. The metagenomic analysis showed the dominance of mesophilic microbiota in cool and temperate sediments, indicating that the downcore decreases of cell abundance and DNA yields are mainly caused by environmental selection against these microbes.

**Figure 3.**
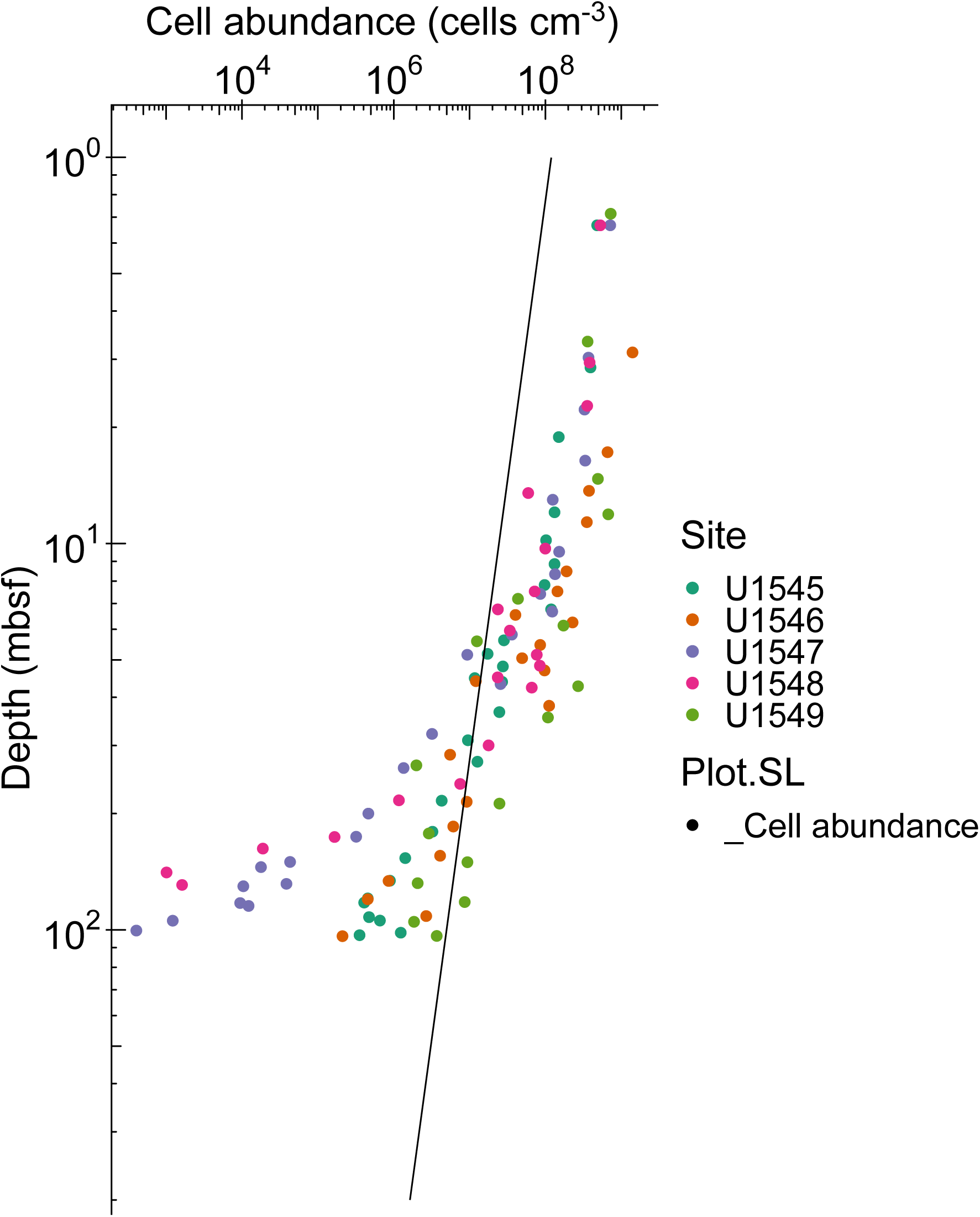
Direct cell counts with SYBR Green I as DNA-staining fluorophore in Guaymas sediment samples from all eight drilling sites. The data pints correspond to samples used for 16S rRNA gene and functional gene surveys of bacteria and archaea (Table 2). Ringvent cell counts cross the Cragg line at 50 mbsf, the other cell counts between ca. 70-100 mbsf.

We note that the depth range horizon for successful metagenomic library construction, sequencing, and recovery of Metagenome-Assembled Genomes (MAGs) is very different from cell count limits. The Expedition 385 summary chapter includes shipboard cell counts that reach 10^5^ to 10^6^ cells/cm^3^ (Teske et al., 2021a). In general, samples with these concentrations yield sufficient DNA for metagenomic sequencing and MAG analysis. Post-cruise counts using automated image acquisition and high-throughput counts in the lab will be much more sensitive, and will allow more precise quantification of subsurface cell density in Guaymas Basin down to 10^2^ to 10^3^ cells per cm^3^ (Morono et al., 2022). Obviously, these sparser deep subsurface microbial communities also contain intracellular DNA that should in principle be amenable to sequencing; yet bridging the sensitivity gap between metagenomics (limited by DNA yield) and cell counts (limited by high-throughput microscopic image processing) remains a challenge. So far, extremely deep and hot samples where cell counts and activity measurements suggest persistent microbial life remain inaccessible by sequencing (Heuer et al., 2020; Beulig et al., 2022).

### Comparison to shallow hydrothermal sediments

Near-surface densities of between 10^9^ and 10^10^ cells/cm^3^ are found in hydrothermal sediments of Guaymas Basin. DNA concentrations in surficial hydrothermal sediments range from 6 µg/cm^3^ (Hinkle et al., 2024) to 10 µg/cm^3^ (Engelen et al., 2021). Based on the average DNA content of 1 to 2 femtograms (10^-15^g) DNA per cell, these DNA concentrations translate into cell densities of 3 to 10 x 10^9^ cells/cm^3^; these numbers match the range of epifluorescence cell count maxima in surficial sediments of Guaymas Basin that range from 1 to 4 x 10^9^ cells/cm^3^ (Meyer et al., 2013). Given high cell numbers and DNA concentrations, metagenomic sequencing of microbial communities in surficial sediments is not limited by DNA availability, and yields highly diverse bacteria and archaeal communities with novel lineages (Dombrowski et al., 2017, 2018; Seitz et al., 2019; Eme et al., 2023).

Consistent with the deep Guaymas Basin subsurface, downcore DNA yields in near-surface hydrothermal sediments of Guaymas Basin decrease by several orders of magnitude, but on scales of centimeters instead of tens or hundreds of meters. Mutually independent studies show that these downcore decreases in DNA yield are persistent, regardless of different DNA extraction protocols and quantification units. DNA yields declined over one order of magnitude from 10 µg DNA/g to below 1 µg DNA/g sediment within the top 3 centimeters of a hot hydrothermal sediment core, using the Qiagen power soil extraction kit and protocol (Engelen et al., 2021). DNA yields declined more than two orders of magnitude from 6 µg/cm^3^ to below 60 ng/cm^3^ within the top 20 centimeters of hydrothermal sediment cores (Table 4 and Figure 4), using a manual extraction protocol that combines freeze-thawing, proteinase K digest, and chloroform-isoamyl alcohol extraction (modified from Zhou et al., 1996). We note that these surficial hydrothermal sediments are very liquid, which minimizes the difference between volume-based and weight-based DNA yield.

**Figure 4.**
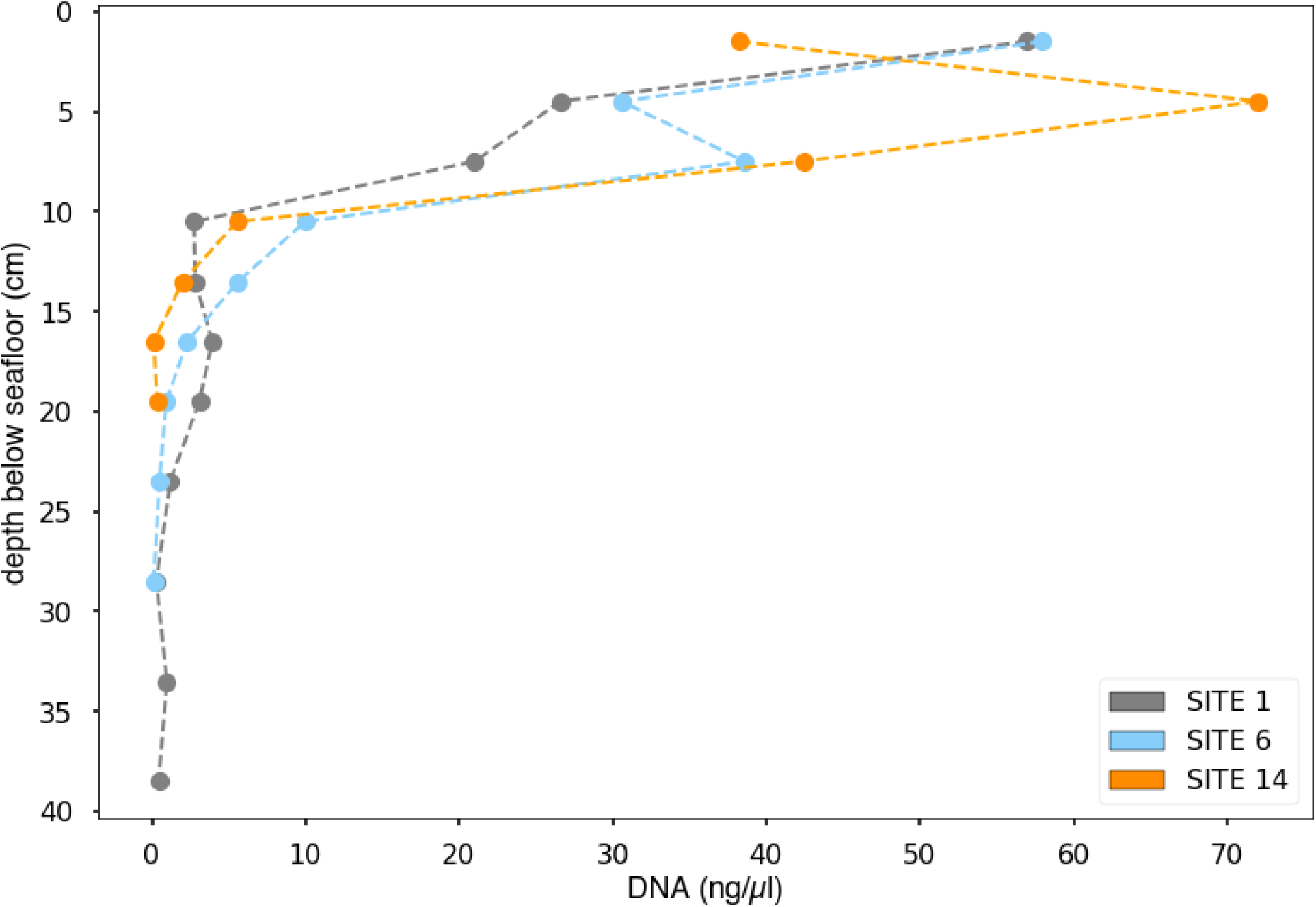
DNA concentration profiles for surficial hydrothermal sediments (Alvin Dive 4872, Dec 24, 2016), extracted manually by a combined freeze-thawing, enzymatic lysis and organic extraction protocol (Zhou et al., 1996) and cleaned up using Amicon purification columns (Table 4). Core 1 was collected in from warm sediment without microbial mat cover, core 6 in hot hydrothermal sediment covered with a white *Beggiatoaceae* mat, and core 14 in very hot hydrothermal sediment covered with an orange *Beggiatoaceae* mat. For final DNA concentrations per mg sediment, the extract concentrations are multiplied by factor 100.

**Table 4.** DNA yields from hydrothermal sediment cores, extracted by J.E. Hinkle and A. Teske. along a gradient from warm sediment without microbial mat cover (core 4872-1) towards the edge of the microbial mat with higher temperatures and white filamentous *Beggiatoaceae* (Core 4872-6), and the hot central mat area with orange *Beggiatoaceae* (core 4872-14) (Alvin Dive 4872, Dec. 24, 2016). DNA concentrations were measured by Qubit (Ruff Laboratory, Marine Biological Laboratory). In-situ temperature measurements via the *Alvin* heatflow probe refer to 0, 10, 20, 30, 40 and 50 cm sediment depth. n.d.: no data; b.d.: below detection limit.

| Alvin core | Depth (cm) | Temp. (°C) | ng DNA per cm<sup>3</sup><br>wet sediment |
| --- | --- | --- | --- |
| 4872-1 | 0-3 | 3 | 5700 |
| 4872-1 | 3-6 | n.d. | 2660 |
| 4872-1 | 6-9 | n.d. | 2100 |
| 4872-1 | 9-12 | 8 | 275 |
| 4872-1 | 12-15 | n.d. | 284 |
| 4872-1 | 15-18 | n.d. | 392 |
| 4872-1 | 18-21 | 11.4 | 310 |
| 4872-1 | 21-26 | n.d. | 120 |
| 4872-1 | 26-31 | 14.6 | 30 |
| 4872-1 | 31-36 | n.d. | 98 |
| 4872-1 | 36-41 | 19.1 | 45 |
| 4872-1 | 41-46 | n.d. | b.d. |
| 4872-1 | 46-51 | 24.8 | b.d. |
| 4872-6 | 0-3 | 3 | 5800 |
| 4872-6 | 3-6 | n.d. | 3060 |
| 4872-6 | 6-9 | n.d. | 3860 |
| 4872-6 | 9-12 | 33.8 | 1000 |
| 4872-6 | 12-15 | n.d. | 562 |
| 4872-6 | 15-18 | n.d. | 232 |
| 4872-6 | 18-21 | 50.9 | 93 |
| 4872-6 | 21-26 | n.d. | 50 |
| 4872-6 | 26-31 | 65.1 | 12 |
| 4872-6 | 31-36 | n.d. | b.d. |
| 4872-6 | 40 | 76.8 | n.d. |
| 4872-6 | 50 | 90.2 | n.d. |
| 4872-14 | 0-3 | 3 | 3820 |
| 4872-14 | 3-6 | n.d. | 7200 |
| 4872-14 | 6-9 | n.d. | 4240 |
| 4872-14 | 9-12 | 83.4 | 556 |
| 4872-14 | 12-15 | n.d. | 202 |
| 4872-14 | 15-18 | n.d. | 13 |
| 4872-14 | 18-21 | 101.8 | 34 |
| 4872-14 | 21-26 | n.d. | b.d. |
| 4872-14 | 26-31 | 105.6 | b.d. |
| 4872-14 | 31-36 | n.d. | b.d. |
| 4872-14 | 36-41 | 108.1 | b.d. |
| 4872-14 | 50 | 102.7 | b.d. |

This strong compression of high DNA yields and microbial populations towards the sediment-water interface is commonly ascribed to increasingly extreme temperatures and biogeochemical conditions in hydrothermally active shallow sediments. These results show some qualitative similarities to the Expedition 385 results, insofar as hydrothermal stress factors appear to compress DNA yield and microbial cell numbers towards the upper subsurface sediments of Guaymas Basin, and depress DNA yield and microbial cell numbers in the subsurface. While the impacts of temperature stress and energy limitation apply to both shallow and deep sediments, the conditions and hydrothermal settings are certainly different. The fluidized surficial hydrothermal sediments that are permeated by pulsating, extremely hot (> 80°C) and highly reducing fluids (McKay et al., 2016; Teske et al., 2016; Su et al., 2023) are distinct from consolidated deeper subsurface sediments characterized by stable thermal and geochemical gradients. Interestingly, the microbial communities of dynamic near-surface hydrothermal sediments and of hydrothermally influenced subsurface sediments are different; the former are dominated by hydrothermal vent and hot spring taxa, whereas the latter share many microbial groups with the global marine sedimentary biosphere (Lagostina et al., 2021). We speculate that microbial communities in hot, hydrothermally active surficial sediments (where numerous carbon sources, nutrients and redox pairs coexist) are less limited by energy and substrate availability than by extreme temperature stress; in contrast, relatively moderate temperatures that are measured in IODP-drilled subsurface sediments have a disproportionally greater impact on the energy-limited microbial deep biosphere. These differences in selection factors may ultimately result in distinct microbial communities.

### Strategies to reach beyond the DNA event horizon

Obtaining sufficient DNA from sediments with lower cell numbers requires further methodological improvements, for example by perhaps separating cells from sediments before cell rupture and DNA extraction to prevent DNA sticking to hydrocarbon-coated sediment particles (Lappé and Kallmeyer, 2011). Refined methods for cell separation from sediments and subsequent concentration have been reviewed and discussed in detail in the context of cell counting (Morono 2023), and will ultimately lead to increased sequencing sensitivity as well.

Bulk sediment DNA isolation procedures can be scaled up using multiple parallel extractions followed by pooling of the extracted DNA, for example by changing from protocol A (1 or 3 parallel extractions per sample) to protocol B (10 parallel extractions per sample). As a caveat, in the course of pooling and concentrating multiple samples, laboratory or reagent contaminants that are present at low levels would also be concentrated and collected. This unintended side effect may have resulted in Actinobacterial and certain Gammaproteobacterial contaminants that appear increasingly in metagenomic sequencing and MAG annotation for low-DNA samples (Mara et al. 2023b). To reduce the need for multiple parallel extractions of small volumes, sediment washing steps similar to those used for RNA isolation could be applied to DNA extraction procedures, to scale up extraction volumes without accumulating inhibitor substances or process-derived contaminants in the final extracts.

With the advent of advanced DNA sequencing technologies, and the laboratory methods to support them, sequencing projects targeting recalcitrant and extreme samples now have a significantly higher rate of success (i.e., Tighe et al. 2017). Reactive and corrosive compounds can co-elute with nucleic acids during typical purification methods, these compounds often negatively impact downstream procedures. Further, organisms that survive in these environments have evolved cellular mechanisms to protect their genomes for gene expression and reproduction. Interestingly, these evolved cellular mechanisms that protect DNA from environmental damage can make the DNA more difficult to extract and to analyze, often inhibiting downstream enzymatic reactions or requiring harsh extraction techniques that results in highly fragmented DNA. New methods and products have been developed that can more effectively remove ‘protective’ compounds and reactive/corrosive compounds. Industry vendors that have stepped into this space with products include Zymo Research (zymoresearch.com), Qiagen (Qiagen.com), Omega Bio-Tek (omegabiotek.com), among others. Home brew methods that have been developed are as simple as performing serial dilutions to reduce inhibitory compounds, or more complex as with newly formulated reagents and solutions. The ability to overcome these challenges has led to a surge in high quality data from previously inaccessible sample materials (Bojanova et al., 2023, Mara et al., 2023).

Computational methods may also expand the limits of useful metagenomic surveys beyond the currently proposed DNA event horizon despite potential contaminant loads. Taxonomy-based identification of contaminants in a metagenomic sequencing datasets may help eliminate unwanted biological information at the short-read, contig assembly, and/or open reading frame (protein prediction) stage. Generating large numbers of contigs that are taxonomically assigned to known contaminants can improve metagenomic assembly quality, by culling the assembly input. Alternatively, independent taxonomic assignments for all proteins predicted in metagenomic contigs can serve as another contaminant culling step with the potential to increase the number of MAGs or the percentage of complete/near complete MAGs from a given dataset (https://github.com/Arkadiy-Garber/Taxonsluice). To extract information from KEGG modules for incomplete bacterial MAGs, MetaPathPredict (Geller-McGrath et al., 2022) can generate predictions for the presence or absence of KEGG modules within gene annotations of bacterial MAGs; we are currently testing this approach for the Guaymas Deep Biosphere (Mara et al. 2023b).

Funding. This study was supported by NSF Grant OCE-2046799 to VE, PM, and AT; by NSF grant OCE-1829903 to VE, PM, and AT; by NASA Exobiology grant APP-0244-001 to AT; by NSF grant OCE-0939564 to D.B., and by JSPS KAKENHI Grants JP19H00730 and JP23H00154 to YM. Expedition 385 participants were aided by IODP cruise and post-cruise support.

## Acknowledgments

We thank all IODP expedition 385 scientists, technicians, drillers and crew for making sample recovery, and by proxy, this research project possible. We gratefully acknowledge the shipboard curatorial team who kept the sediment samples and metadata organized and well-catalogued. We thank the Amend Lab for hosting G. Ramírez for the first rounds of DNA extractions before the Pandemic shutdown. We thank Eleanor Greene and Hannah Vanderscheuren (Ruff Lab, MBL) for clean-up and quantification of the DNA in Figure 4.

